# Thermal cycling transcription boosts RNA production

**DOI:** 10.1101/2025.01.18.633700

**Authors:** Yuxuan Liu, Lu Shi, Juergen Brosius, Dingding Mo

## Abstract

Efficient RNA production is essential for therapeutic applications. Currently, mRNA is generated by *in vitro* transcription with RNA polymerases at a constant temperature. The success of these transcriptions depends on various factors including sequence composition and size. Sometimes, especially for long RNAs, achieving efficient transcript yields are challenging.

We developed a novel approach by incorporating a high-temperature denaturation step. Our protocol includes the following two steps of thermal cycling RNA transcription: the initial enzyme-driven RNA transcription step is done at RNA polymerase active temperature (37 °C); then the nucleic acids melting step is carried out at elevated temperatures (55–70°C); after the reaction is cooled to the temperature of optimal polymerase activity, the first step is repeated.

Denaturation enables the generation of optimized, more stable mRNAs, which alleviates the mRNA instability that often occurs during RNA-based therapeutic development. For example, this new thermal cycling RNA transcription approach can employ high GC content, thus increasing RNA stability.

For long and difficult templates, the combination of sequence optimization by LinearDesign algorithm in conjunction with transcription via thermal cycling significantly improved mRNA production. Thermal cycling transcription dramatically increased the efficiency of most RNA products while reducing costs, facilitating RNA-based therapeutic development.

## Introduction

mRNA-based therapies are developing at a rapid pace. However, apart from use in vaccines, several key obstacles are encountered during the application of mRNA for medical use. For example, compared to mRNA vaccines, about a 1,000-fold higher protein level is required for therapeutic threshold (1). Other problems include RNA instability, impeding production, storage and mRNA delivery. Recent studies indicate that increased stability and translatability can be achieved by increasing the guanosine/cytidine (G/C) ratio and stabilizing RNA secondary structures; applying such techniques as LinearDesign algorithm optimization to mRNAs can achieve this (2).

The current standard for mRNA production is *in vitro* transcription (IVT); significantly increasing IVT efficiency will dramatically reduce RNA production costs. IVT efficiency, especially the efficiency of T7 RNA polymerase, an extensively-used *in vitro* transcription enzyme, has been well studied (3-5). However, T7 RNA polymerase is not always successful in transcribing every individual DNA template (6-9). Sequence composition and mRNA sizes significantly affect transcription efficiency. For the transcription of some highly structured mRNAs, T7 RNA polymerase can be slowed down or even blocked by G/C rich sequences and very stable hairpin-shaped secondary structures, which also may resemble transcription terminators and cause internal transcription termination, leading to low yields of run-off mRNA production or to the accumulation of prematurely truncated transcripts (7-9). Thus, establishing a universal technology that produces sequence optimized mRNA that can encode any target protein would greatly improve the modality and versatility of mRNA medicine.

Here we developed a robust *in vitro* transcription technology by alternating the incubation temperatures during transcription between the optimal temperature for RNA polymerase activity and high temperature for the purpose of denaturation. Our innovation dramatically improved RNA production.

## Materials and Methods

### DNA template preparation

The mRNA expression plasmid was synthesized at Genscript. The expression cassette included the T7 transcription promoter, a 5’UTR, the mCherry coding sequence, a3’UTR and 97nt long poly(A) stretch. The expression cassette was PCR amplified, *Bsp*QI digested and phenol-chloroform purified as transcription template. The mCherry mRNA sequence was optimized using LinearDesign algorithm (2). The two alternative AUG translational start codons and the UGA stop codon are indicated by bold letters.

#### Sequence of mCherry mRNA

GGGAUA**AUG**GUACCCUGCAGGCUAGCAUUCUUCUGGUCCCCACAGACUCAGAGAGAACCCGCCACC<u>**AUG**GUGAGCAAGGGGGAGGAGGAUAACAUG GCCAUUAUCAAGGAGUUCAUGCGCUUCAAGGUGCAUAUGGAGGGCUCCGUUAAUGGCCAUGAGUUCGAGAUCGAGGGCGAAGGGGAGGGGCGCCCC UAUGAGGGCACUCAGACUGCGAAGCUGAAGGUCACCAAGGGGGGGCCCCUCCCCUUCGCCUGGGAUAUCCUCUCCCCCCAGUUCAUGUACGGCUCGA AGGCAUACGUCAAGCAUCCUGCGGACAUCCCCGACUACUUGAAGCUCUCCUUCCCGGAGGGCUUCAAGUGGGAGCGGGUGAUGAACUUCGAGGACGG UGGAGUGGUGACCGUCACGCAGGACUCGUCGCUCCAGGAUGGUGAGUUCAUCUACAAGGUGAAGCUCCGAGGAACGAACUUCCCAUCCGAUGGUCCU GUGAUGCAGAAGAAGACCAUGGGAUGGGAAGCGUCCUCGGAGCGUAUGUACCCGGAGGACGGGGCCUUGAAGGGGGAGAUCAAGCAGCGUCUUAAG CUCAAGGACGGUGGGCAUUAUGACGCUGAGGUCAAGACCACCUACAAG GCGAAGAAGCCCGUCCAGCUUCCGGGUGCAUACAACGUGAAUAUCAAGCUUGAUAUCACGUCUCACAAUGAGGACUACACCAUCGUGGAGCAGUAC GAGCGUGCGGAGGGUCGCCACUCCACCGGAGGUAUGGACGAGCUGUAC AAG**UGA**</u>AAACGAAUUCGCGGCCGCAUGCAUGUCGACUAAUAGUAGGCU GGAGCCUCGGUGGCCAUGCUUCUUGCCCCUUGGGCCUCCCCCCAGCCCCUCCUCCCCUUCCUGCACCCGUACCCCCGUGGUCUUUGAAUAAAGUCUGAGUGGGCGGCACGUCUCUUAACUAACUAAGGAUCCCGUCUCUUAACUAACUAAACUAGUAAAAAAAAAAAAAAAAAAAAAAAAAAAAAAAAAAAAAAAAAAAAAAAAAAAAAAAAAAAAAAAAAAAAAAAAAAAAAAAAAAAAA AAAAAAAAAAAA

### RNA Transcription and 5’ capping

RNA Transcription was performed under the following conditions: 100 mM Hepes-K (pH 7.9), 24 mM MgCl_2_, 30 mM DTT, 4 mM Spermidine, 4 mM rNTP each, 200 U/ml RNase inhibitor (10777019, Invitrogen), 10 U/ml thermostable inorganic pyrophosphatase (M0296L, NEB), 0.16 mg/ml T7 RNA polymerase (purified via Ni-NTA and heparin affinity chromatography), 50 µg/ml DNA template with T7 promoter. For mRNA synthesis, N1-Me-Pseudo UTP (HBP002501, Hzymes) was used instead of UTP. RNA transcription was performed either at constant temperature, such as 37 °C or 42 °C, or at alternating temperatures. Thermal cycling RNA transcription: 20 times thermal cycling of transcription at 37 °C or 42 °C for 3 minutes and denaturation at high temperature (55 °C–75 °C) for 1 minute. mRNA was capped with vaccinia virus capping enzyme (HBP000611, Hzymes).

### RNA purification

RNA transcripts were digested with 80 U/ml DNase I (04716728001, Roche) to remove DNA templates followed by phenol-chloroform extraction to remove proteins. Finally, RNA was purified from agarose gel electrophoresis and electroelution.

### Cell culture and RNA transfection

HEK293T and A549 cells were cultured in DMEM medium (C11995500CP, Gibco) supplemented with 10% fetal bovine serum (10091148, Gibco), 100 U/ml penicillin and 100 U/ml streptomycin (15140122, Gibco) at 37 °C in 5% (v/v) CO_2_. 3 µg mRNA was transfected with 5 µl Lipofectamine™ MessengerMAX (LMRNA003, Invitrogen) in 6 well plates.

## Results

### Thermal cycling boosts mCherry mRNA production

Conventional RNA transcription was performed at constant temperatures, such as 37 °C. We designed an mCherry mRNA *in vitro* transcription system. The DNA template includes the T7 promoter, 5’ UTR (from human hemoglobin subunit alpha 2), mCherry open reading frame (sequence was optimized using LinearDesign algorithm (2)), 3’ UTR (from human hemoglobin subunit alpha 1) and poly(A) tail (97nt) (Fig. 1A). DNA template was amplified from the plasmid construct by PCR and purified by phenol-chloroform extraction. The conventional *in vitro* RNA transcription was performed as described in Methods section. As shown in Fig. 1B, mCherry mRNA transcription at 37 °C generated moderate levels of mRNA.

**Fig. 1.**
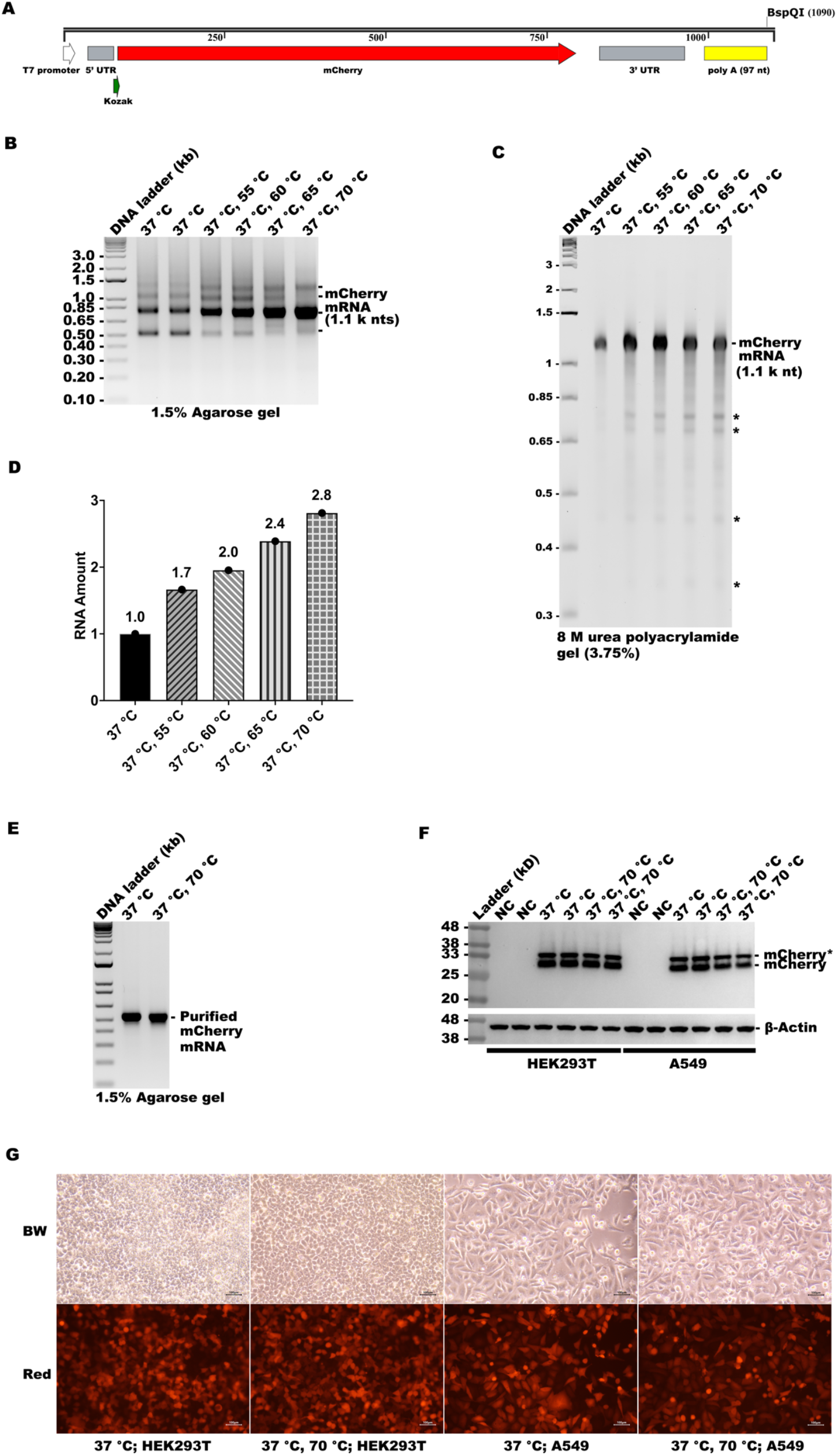
Thermal cycling transcription boosts mCherry mRNA production. **A**. Schematic of the DNA template for mCherry mRNA production (the mCherry ORF was optimized by LinearDesign program). 5’ UTR, 5’ untranslated region of human hemoglobin subunit alpha 2 (HBA2); 3’ UTR, 3’ untranslated region of human hemoglobin subunit alpha (HBA1); poly(A) (97nt), the 97nt poly(A) tail. **B**. Agarose gel (1.5%) electrophoresis of mCherry mRNA transcription products (phenol-chloroform purified); ‘‘37 °C’’, RNA transcription at 37 °C for 1 hour; ‘‘37 °C, 55 °C’’, 20 times thermal cycling of transcription at 37 °C for 3 minutes and denaturing at 55 °C for 1 minute; ‘‘37 °C, 60 °C’’, 20 times thermal cycling of transcription at 37 °C for 3 minutes and denaturing at 60 °C for 1 minute; ‘‘37 °C, 65 °C’’, 20 times thermal cycling of transcription at 37 °C for 3 minutes and denaturing at 65 °C for 1 minute; ‘‘37 °C, 70 °C’’, 20 times thermal cycling of transcription at 37 °C for 3 minutes and denaturing at 70 °C for 1 minute. DNA ladder was used as size marker. **C**. Polyacrylamide gel (3.75%) with 8 M urea electrophoresis of phenol-chloroform purified mCherry mRNA transcription products; -, mCherry mRNA (1.100nt); *, by-products. **D**. quantification of the mCherry mRNA amounts in B. **E**. Agarose gel (1.5%) electrophoresis of purified mCherry mRNA transcribed in A and then capped with Vaccinia Virus Capping Enzyme. **F**. Western blot analysis of the HEK293T and A549 cells transfected with the purified and 5’ capped mCherry mRNA of two transcription protocols (‘‘37 °C’’, ‘‘37 °C, 70 °C’’); -, mCherry protein; *, mCherry protein with the N-terminal 2.4 kDa extension initiating from the upstream AUG in the 5’ UTR region of mCherry mRNA. NC, non-infected negative control cells; **G**. Fluorescence of mCherry in HEK293T and A549 cells transfected with the purified mCherry mRNA of two transcription protocols (‘‘37 °C’’, ‘‘37 °C, 70 °C’’); BW, black and white view of cells; Red, red fluorescence of mCherry.

Subsequently, we performed thermal cycling RNA transcription which is described in detail in the methods section. Agarose gel electrophoresis showed that the addition of denaturing step at 55 °C, 60 °C, 65 °C, or 70 °C significantly increased mRNA yields concomitantly rising with the denaturation temperature (Fig. 1B). Quantification of Fig.1B showed that thermal cycling increased RNA production from 1.7-fold to 2.8-fold (Fig. 1D). To verify the mRNA length of thermal cycling transcription, we performed 8 M urea-polyacrylamide denaturing gel (3.75%) electrophoresis of the synthesis products. As shown in Fig.1C, thermal cycling transcription produced mCherry mRNA run-off product of the same size as in the constant 37 °C transcription, but once more, with significantly increased yields. The short by-products, presumably due to premature termination, were only moderately increased (Fig.1C). The denaturing gel also indicates that the additional bands detected in the native agarose gel were possibly secondary structure variants with different migration properties (Fig. 1B). After treatment with vaccinia virus capping enzyme, mCherry mRNAs was purified by agarose gel electrophoresis and electroelution. The purified mCherry mRNAs produced at constant 37 °C or at thermal cycling transcription (‘‘37 °C, 70 °C’’) were once more analyzed by agarose gel electrophoresis, showing identical migration patterns (Fig.1E). Furthermore, we transfected these two purified mRNAs into HEK293T and A549 cells. After 18-hour transfection, both expressed similar levels of mCherry protein in HEK293T and A549 cells, demonstrated by Western blot analysis of mCherry protein with anti-mCherry antibody (Fig.1F) and by direct mCherry red fluorescence in the cell cultures (Fig.1G). During template design, an additional in-frame AUG start codon was inadvertently introduced into the 5’ UTR. Despite the absence of an obvious Kozak sequence (10), it apparently also served as translation initiation codon leading to an additional N-terminal sequence of 20 amino acids, hence the observed double band. All evidence indicated that mCherry mRNA templates generated by thermal cycling are equally efficient in their capacities to be translated into a polypeptide compared to those transcribed at constant 37 °.

### Thermal cycling boosts the transcription of other LinearDesign algorithm optimized mRNAs

As thermal cycling transcription significantly improved mCherry mRNA production, we wished to explore whether this approach could generally be applied to the synthesis of other mRNAs. The open reading frames of three mRNAs for proteins used in genome engineering (ISAam1 (11), SpuFz1-V2 (12) and Cas9 (13)) were inserted into the mRNA expression cassette (replacing mCherry), including two mRNA variants for all three mRNAs, one was the original mRNA sequence and the other variant was optimized by LinearDesign algorithm. Six DNA templates were prepared similarly to the mCherry DNA template by PCR.

Surprisingly, mRNA transcription of the original 1661nt ISAam1 sequence produced somewhat more intact mRNA at constant 37 °C than during thermal cycling (‘‘37 °C, 70 °C’’) transcription (Fig. 2A). This result is possibly explained by the instability of the original ISAam1 mRNA sequence during the denaturing step at 70 °C. In fact, the LinearDesign algorithm optimized ISAam1 mRNA had a better yield than the natural mRNA generated at constant 37 °C (Fig. 2A); thermal cycling somewhat increased its yield (Fig. 2A).

**Fig. 2.**
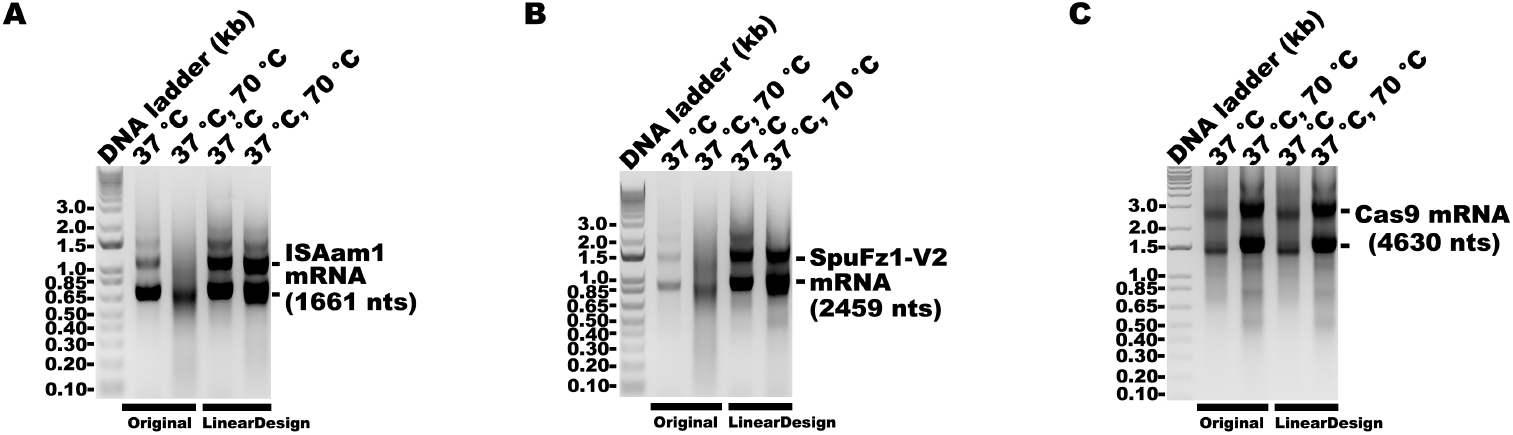
Thermal cycling transcription boosts other mRNA productions. **A, B, C**. Agarose gel (1.0%) electrophoresis of ISAam1, SpuFz1-V2, and Cas9 mRNA transcription products (phenol-chloroform purified); ‘‘37 °C’’, RNA transcription at 37 °C for 1 hour; ‘‘37 °C, 70 °C’’, 20 times thermal cycling of transcription at 37 °C for 3 minutes and denature at 70 °C for 1 minute; Original, the original ISAam1/SpuFz1-V2/ Cas9 sequence; LinearDesign, optimized ISAam1/SpuFz1-V2/ Cas9 sequence; DNA ladder was used size marker.

Transcription of the original 2459nt SpuFz1-V2 mRNA at both constant 37 °C and thermal cycling (‘‘37 °C, 70 °C’’) conditions produced only limited mRNA (Fig. 2B), indicating the challenges of this template for transcription. LinearDesign algorithm optimized SpuFz1-V2 improved mRNA transcription output at 37 °C (Fig. 2B). Thermal cycling (‘‘37 °C, 70 °C’’) further boosted its yield (Fig. 2B).

Finally, the original 4630nt Cas9 mRNA expression cassette produced very limited mRNA output at constant 37 °C (Fig. 2C). Thermal cycling (‘‘37 °C, 70 °C’’) significantly improved the mRNA yield (Fig. 2C). Surprisingly, LinearDesign algorithm optimized Cas9 mRNA also had a lower level of transcription at constant 37 °C. But thermal cycling transcription (‘‘37 °C, 70 °C’’) once again significantly improved the mRNA output (Fig. 2C), however, without much difference between the natural and improved form. In conclusion, thermal cycling boosts the transcription of LinearDesign algorithm optimized mRNAs.

## Discussion

We add We developed a new strategy to overcome some of the transcription obstacles of *in vitro* mRNA synthesis. Thermal cycling, widely-used in PCR procedures, was applied to mRNA transcription. ed a denaturing step between mRNA transcription conditions. Surprisingly, T7 RNA polymerase tolerated heating in the denaturation step up to 75 °C (data not shown), omitting the requirement of adding fresh enzyme after each cycle. In most cases, the denaturing step effectively boosted transcription, even of large transcripts (such as the 1.100nt mCherry mRNA and the 4.600nt Cas9 mRNA).

The denaturing step may reduce the intrinsic RNA secondary structure, thus minimizing the T7 terminator resembling structures mediated transcription terminations. The denaturing step may also facilitate the diffusion of T7 RNA polymerase back to the promoter region of DNA template for reinitiation of transcription (14).

For the transcription of difficult templates (such as 2459nt SpuFz1-V2 mRNA and 4630nt Cas9 mRNA), the combination of sequence optimization by LinearDesign algorithm and transcription by thermal cycling jointly and in some cases both significantly improved mRNA yields, thus representing a universal mRNA synthesis approach for any individual protein, further enhancing the therapeutic options and versatilities of mRNA therapeutics.

## Materials and data availability

Requests for further information and resources should be directed to and will be fulfilled by Dingding Mo.

## Declaration of interests

D.M. is inventor of patent containing data published in this paper. Other authors declare no competing interests.

## Funding

### Authors’ contributions

D.M. designed, conceived and supervised the study. D.M. analysed the results, prepared the figures, and wrote the manuscript. Y.L. performed the experiments. L. S. performed the mRNA sequence optimizations. J.B. participated in data analysis, manuscript editing and revising.

## Acknowledgements

Authors acknowledge the kind support of Prof. Sheng Ding during the study. We also thank Dr. Lin Huang and Dr. Keqiong Ye for T7 RNA polymerase construct and RNA transcription protocol as well as Stephanie Klco-Brosius for editorial advice.

